# Piezo1 activates non-canonical EGFR endocytosis and signaling

**DOI:** 10.1101/2022.05.10.490586

**Authors:** Carlos Pardo-Pastor, Jody Rosenblatt

## Abstract

**Summary:** EGFR-ERK signaling controls cell cycle progression during development, homeostasis, and disease. Both the soluble extracellular ligand, EGF, and mechanical inputs like matrix stiffness, cell adhesion, or stretch activate EGFR-ERK signaling. However, the molecules transducing mechanical forces into EGFR signaling remain unidentified. We previously found that stretch promotes mitosis by mechanically activating the ion channel Piezo1 to trigger ERK signaling. Here, we show that Piezo1 provides the missing link to mechanical EGFR-ERK activation. Both EGF ligand and mechanical or agonist activation of Piezo1 trigger clathrin-mediated endocytosis of EGFR and ERK activation, established markers of EGFR signaling. However, while EGF stimulation requires canonical EGFR tyrosine autophosphorylation, EGFR activation by Piezo1 instead requires Src-p38 kinase-dependent serine EGFR phosphorylation. Additionally, in contrast to the homeostatic signaling downstream EGF activation, direct agonist stimulation of Piezo1 promoted cell cycle re-entry and proliferation in mouse airway epithelia *ex vivo* via long-term nuclear accumulation of ERK, AP-1 (FOS and JUN), and YAP, typical of regenerative and malignant signaling. Our results suggest two modes of EGFR signaling: basal EGF-dependent signaling via tyrosine autophosphorylation and mechanically activated Piezo1-dependent signaling via serine phosphorylation, resulting in sustained proliferation, critical to repair, regeneration, and cancer growth. Given the limited success of tyrosine kinase inhibitors in cancer, this new axis may provide a more relevant target for cancer treatment.

## Introduction

Mechanical regulation of cell proliferation, differentiation, and migration was originally described decades ago (*1–3*), but only recently have molecules that integrate mechanics and signaling come to light. The mechanosensitive Piezo ion channels and YAP/TAZ transcriptional regulators are the most noteworthy in this integration (*4–7*). Human Piezo1 and 2 are mechanically activated cation channels, abundant in a wide variety of tissues and cell types (*4*), that mediate extracellular calcium influx in response to cell confinement (*8*), stretch (*9*), shear stress (*10–12*), or matrix stiffness (*13, 14*). Piezo-dependent calcium entry is critical for development, regeneration, and cancer through its roles in cell migration (*8, 13, 15*), matrix remodeling (*13, 16*), cytokinesis (*9*), or stiffness-dependent nuclear translocation of YAP (*13, 14*). We found that Piezo1 promotes epithelial cell extrusion in response to crowding (*17*) and rapid cell division (G_2_/M transition) in response to stretch (*18*). Moreover, stretch also promotes cell exit from quiescence (G_0_/G_1_ transition) and DNA synthesis (G_1_/S transition) (*19*), both requiring Piezo1 (*20*). These studies strengthen the role of mechanical forces and Piezo1 as master regulators of the cell cycle.

Soluble growth factors regulate cell function through membrane receptors, too. <u>E</u>pidermal <u>G</u>rowth <u>F</u>actor <u>R</u>eceptor (EGFR) is a widely expressed Receptor Tyrosine Kinase that autophosphorylates its C-terminal cytoplasmic tyrosine residues after binding extracellular soluble ligands (*21–26*). These modified residues act as docking sites for adaptor proteins containing phosphotyrosine (pY)-binding SH2 domains, such as SHC and GRB2. Both endocytic (AP2, Clathrin) and effector proteins (GEFs-Ras-MAPK, PKB/AKT, PLC, PI3K) bind pY-bound adaptors, and their specific combination and phosphorylation by EGFR triggers receptor endocytosis and downstream signaling pathways, such as RAS-MAPK, that impact cell migration, cycling (cell division, quiescence and DNA synthesis), and malignancy (*21–23, 25, 27–35*). In response to high ligand doses, EGFR becomes downregulated by a ubiquitin-dependent, non-clathrin endocytic mechanism that targets the receptor for destruction (*26, 36–38*).

Importantly, mechanical signals such as cell stretch (*28, 29*), apicobasal compression (*39*), matrix-cell adhesion and cell spreading (*31, 32, 40, 41*), or substrate stiffness (*22, 30, 34*) can also activate EGFR endocytosis and signaling. However, little is known about molecules that transduce mechanical forces into EGFR signaling and which EGFR pathways mechanical activation induces. As we previously found that Piezo1 controls stretch-activation of the MAPK ERK (*18*), a target of EGFR, here, we investigate if Piezo1 provides a molecular link between mechanical and EGFR signaling.

## Results

### Piezo1 triggers clathrin mediated endocytosis of EGFR

To investigate the role of Piezo1 in mechanical activation of EGFR signaling, we exposed cells to shear stress, a previously identified Piezo1-activating mechanical stimulus (10–12), and monitored internalization of EGFR phosphorylated on tyrosine 1173 (pY1173-EGFR), an indicator of active EGFR signaling (22, 40, 42, 43). We chose HeLa cells because they endogenously express Piezo1 and are widely used for studying EGFR (26, 38, 43–45). Serum-starved HeLa cells exposed to shear stress for 15 minutes redistributed pY1173-EGFR from the cell periphery to intracellular puncta that colocalize with the early endosome marker EEA1 (Fig.1A-C, F). To test if shear stress-dependent EGFR signaling requires Piezo1, we knocked down Piezo1 with siRNA, confirming its loss by the suppression of calcium transients in response to the synthetic Piezo1 agonist, Yoda1 (Fig.S1A-C). Importantly, siRNA-mediated knockdown of Piezo1 (siPiezo1) or Clathrin Heavy Chain (siCHC, key endocytic protein (26, 38)), impaired pY1173-EGFR internalization by shear stress, resembling unstimulated siControl cells (Fig.1B, C, F), showing that mechanical activation of EGFR endocytosis requires Piezo1 and clathrin.

**Figure 1.**
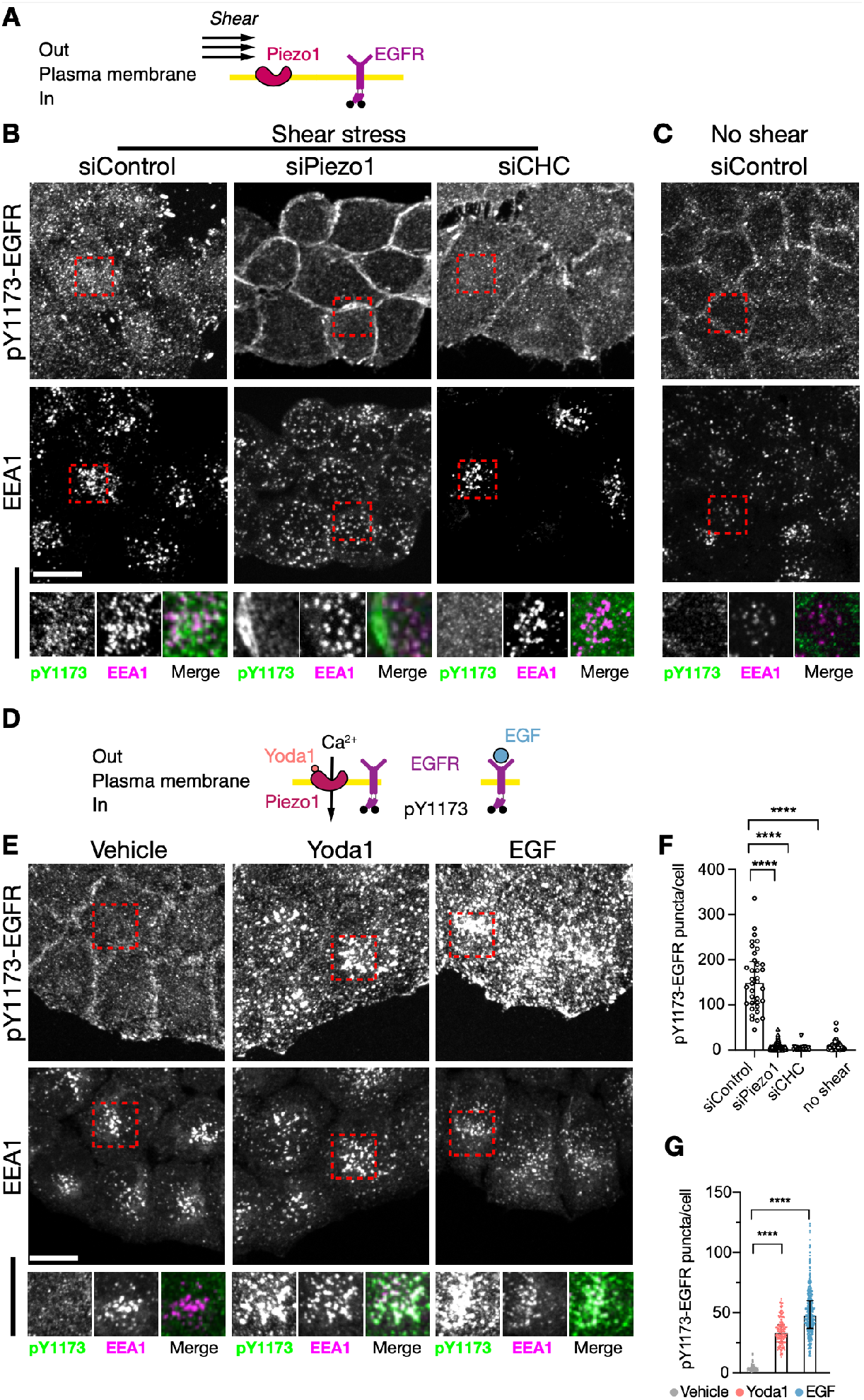
Clathrin mediated endocytosis of EGFR in response to Piezo1 activation. **A, D:** Schematic of the experimental set-ups. **B, C, E:** Representative maximum intensity projections of pY1173-EGFR and EEA1 co-stainings in HeLa cells treated for 15min as indicated. **F, G:** Counts of pY1173-EGFR puncta per cell. Each symbol represents a cell (F, siCtrl: 36, siPiezo1: 161, siCHC: 28, no shear: 44; G, Vehicle: 389, Yoda1: 454, EGF: 925, 4 experiments). Scale bar = 20μm. Error bars = median±interquartile range. **** p< 0.0001 in one-way ANOVA followed by Kruskal-Wallis post-hoc test with Dunn’s correction for multiple comparisons.

To directly activate Piezo1 without triggering other mechanosensitive pathways, we treated serum-starved HeLa cells with the Piezo1 agonist Yoda1 (46). 15-minute incubations with 10 µM Yoda1 or 10 ng/mL EGF (as positive control) redistributed pY1173-EGFR from the cell periphery to intracellular EEA1+ puncta (Fig.1D-F). In addition, siPiezo1 suppressed responses to Yoda1, but not to EGF. Knockdown of Clathrin Light Chain suppressed pY1173-EGFR internalization in all scenarios (Fig.S1D, E). Moreover, Yoda1-induced EGFR internalization was conserved in human lung adenocarcinoma A549 and canine kidney MDCK cells (Fig.S1F), widely used in EGFR studies (47–50), suggesting a conserved mechanism. A phosphorylation-independent EGFR antibody further confirmed EGFR internalization in response to Piezo1 activation (Fig.S1G). Together, these results both demonstrate that mechanical stimuli and Yoda1 trigger Piezo1-dependent EGFR internalization into early endosomes via Clathrin Mediated Endocytosis (CME), a pathway shared with canonical EGFR signaling in response to low EGF doses (26).

### EGFR kinase activity and tyrosine phosphorylation are dispensable for Piezo1-triggered EGFR endocytosis

Both canonical ligand-dependent EGFR phosphorylation and internalization or its transactivation by other transmembrane proteins, calcium signaling, or mechanical forces relies on ligand-receptor interaction and EGFR tyrosine autophosphorylation (*24, 25, 28, 39, 51*). Given that Piezo1 is a transmembrane protein that mediates calcium influx in response to mechanical forces (*4*), we tested if Piezo1-dependent EGFR signaling requires receptor autophosphorylation. While the EGFR kinase inhibitor PD153035 prevented EGFR internalization in response to EGF, as expected (*28, 48*), it did not impact EGFR internalization in response to Yoda1 (Fig.2 A-C). In addition, Yoda1 treatment did not increase pY1068-or pY1173-EGFR, two major autophosphorylation sites (*22, 40, 42, 43, 48*) (Fig.2D, E). Thus, we conclude that Piezo1-initiated EGFR endocytosis does not require canonical tyrosine autophosphorylation.

**Figure 2.**
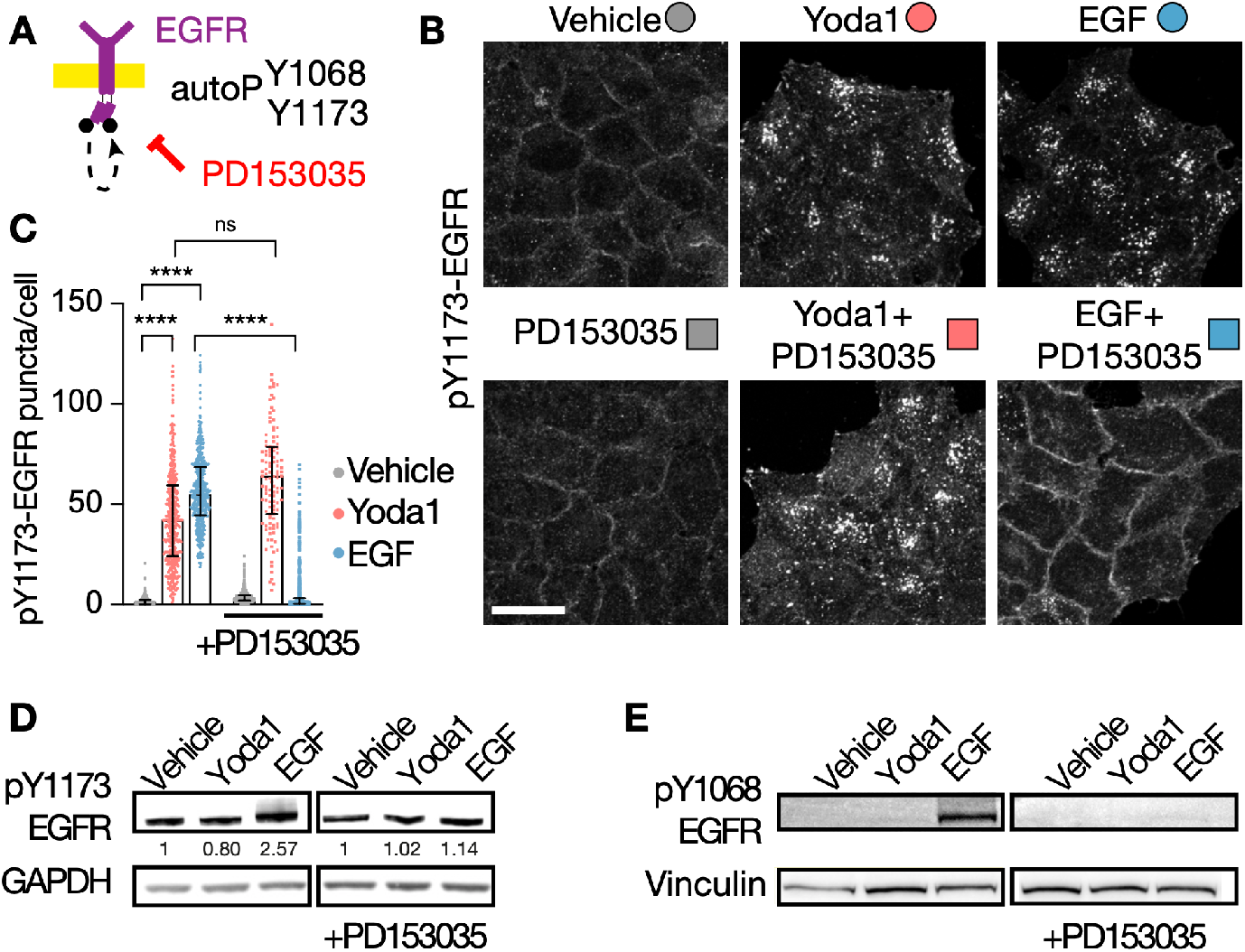
EGFR kinase activity and tyrosine phosphorylation are dispensable for Piezo1-triggered EGFR endocytosis. **A:** Schematic of the experimental set-up. **B:** Representative maximum intensity projections of pY1173-EGFR stainings in HeLa cells treated for 15min with vehicle, 10μM Yoda1 or 10ng/mL EGF with or without 30min preincubation with 1 μM PD153035 (EGFR kinase inhibitor). **C:** Counts of pY1173-EGFR puncta per cell after the indicated treatments. Each symbol represents a cell. Vehicle: 2725, Yoda1: 860, EGF: 1766; PD: 2136, Yoda1+PD: 405, EGF+PD: 2410, 6 experiments. **D, E:** Representative western blots of pY1173- and pY1068-EGFR. GAPDH and Vinculin used as loading controls. Scale bar= 20μm. Error bars = median±interquartile range. ns = non-significant, **** p< 0.0001 in one-way ANOVA followed by Kruskal-Wallis post-hoc test with Dunn’s correction for multiple comparisons.

### Non-canonical EGFR serine phosphorylation by an SFK-p38 axis in response to Piezo1 activation

Cell-matrix adhesion regulates EGFR via the Src Family Kinases (SFK) (*32, 40*) and can bypass EGFR kinase activity (*41*), suggesting an alternative signaling axis to regulate EGFR in response to Piezo1 activation. HeLa cell pre-treatment with the SFK inhibitor PP2 suppressed Yoda1-induced pY1173-EGFR internalization but did not affect responses to EGF (Fig.3A-C). This shows that EGFR endocytosis in response to Yoda1 requires SFK activity, unlike EGF-mediated signaling. Although adhesion-dependent EGFR signaling requires SFK-dependent phosphorylation of EGFR on Y1068 (*41*), we find that Yoda1 does not alter pY1068-EGFR levels (Fig.2E). Therefore, we investigated other non-canonical mechanisms that might control EGFR endocytosis by Piezo1.

**Figure 3.**
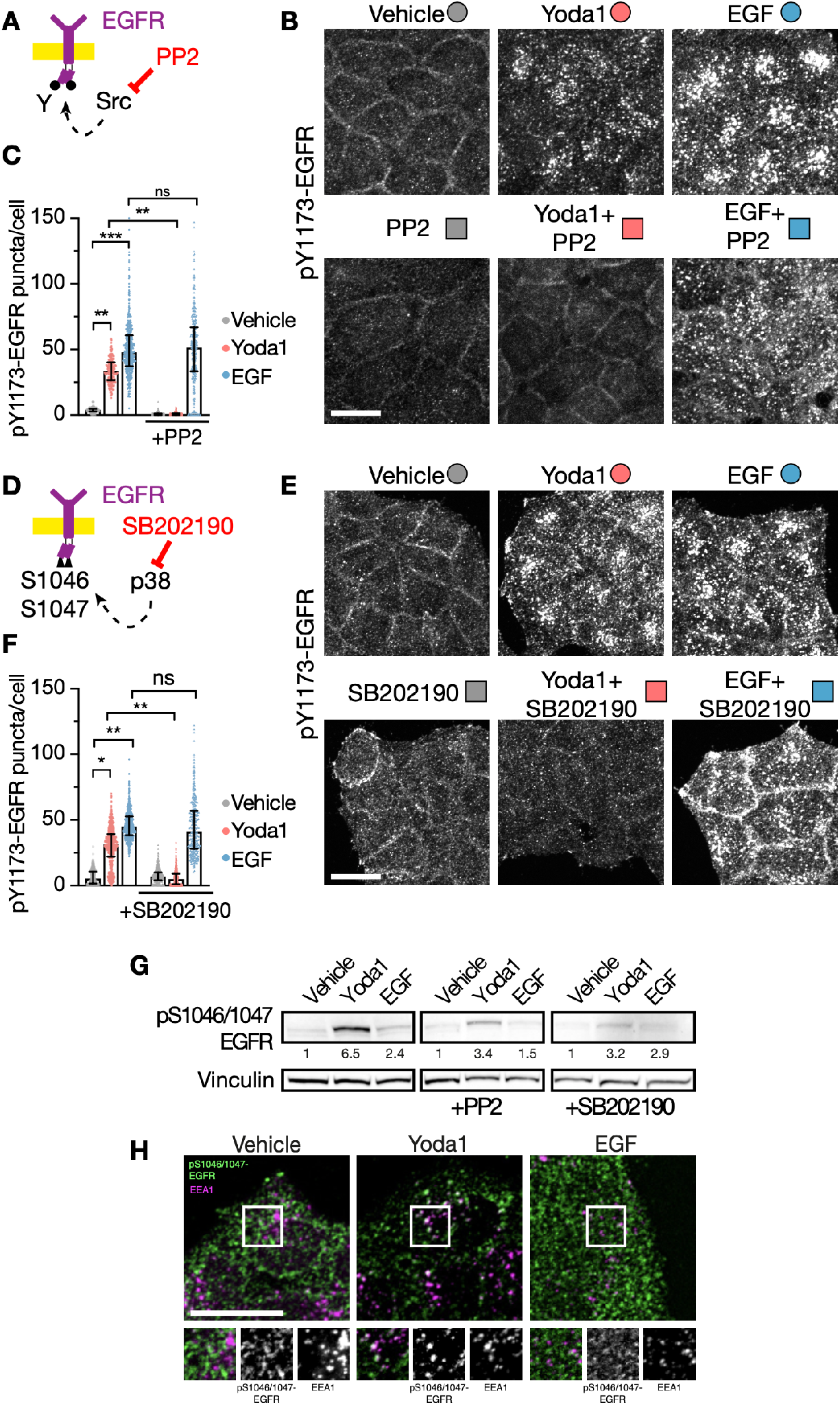
Non-canonical EGFR serine phosphorylation by an SFK-p38 axis in response to Piezo1 activation. **A, D:** Schematic of the experimental set-ups. **B, E, H:** Representative maximum intensity projections of pY1173-EGFR (B, E) or pSer1046/1047-EGFR (H) stainings in HeLa cells treated for 15min with vehicle, 10μM Yoda1 or 10ng/mL EGF with or without 200nM PP2 (SFK inhibitor, B), or 10 μM SB202190 (p38 inhibitor, E). **C, F:** Counts of pY1173-EGFR puncta per cell after the indicated treatments. Each symbol represents a cell. Vehicle: 1143, Yoda1: 717, EGF: 32, PP2: 1694, Yoda1+PP2: 887, EGF+PP2: 829, 6 experiments (C); Vehicle: 867, Yoda1: 1125, EGF: 1544, SB: 948, Yoda1+SB: 1358, EGF+SB: 1016, 3 experiments (F). **G:** Representative western blots of pS1046/1047-EGFR. Vinculin used as loading control. Scale bar= 20μm. Error bars = median±interquartile range. ns = non-significant, * p<0.05, ** p<0.01, *** p<0.001 one-way ANOVA followed by Kruskal-Wallis post-hoc test with Dunn’s correction for multiple comparisons.

Because Piezo1 can activate p38 (*52*), we tested if p38 kinase mediates Piezo1-dependent EGFR endocytosis. p38 MAPK has been previously found to phosphorylate EGFR serine residues 1046 and 1047 (pS1046/1047-EGFR) in response to cellular stresses, which in turn recruits the endocytic machinery (e.g. AP2, clathrin) to internalize EGFR (*44, 47, 48, 53–56*). We find that inhibiting p38 with SB202190 suppresses EGFR endocytosis in response to Yoda1, but not to EGF (Fig.3D-F). Additionally, immunoblots indicate that Yoda1 but not EGF increases pS1046/1047-EGFR in an SFK-dependent fashion (Fig.3G). Accordingly, Yoda1 but not EGF also increased the number of pS1046/1047-EGFR^+^ endosomes (Fig.3H). Thus, Piezo1 activation leads to SFK- and p38-dependent EGFR serine phosphorylation to drive receptor endocytosis, a pathway shared with cell stress (*44, 47, 48, 53–55*).

### Functional EGFR internalization in response to Piezo1 activation leads to different nuclear signals

ERK1/2 MAPK activation is a hallmark of EGFR signaling, with amplitude, duration, and location determining pERK1/2 signaling outcomes (*33, 57*). To investigate how Piezo1 activation of EGFR signaling affects ERK1/2 signaling, we next followed total ERK1/2 activation and localization. Immunoblots indicate that both Yoda1 and EGF activation of EGFR promote similar levels of (active) phosphorylated ERK1/2 (pERK1/2) (Fig.4A, left lanes). While the EGFR kinase inhibitor PD153035 suppressed EGF-dependent pERK1/2, it only partially inhibited phosphorylation by Yoda1 (Fig.4A, center lanes). However, the SFK inhibitor PP2 dampened both Yoda1- and EGF-induced pERK1/2 (Fig.4A, right lanes). Interestingly, pERK1/2 accumulates within nuclei in response to Yoda1 but not EGF treatment (Fig.4B, C). As in previous experiments, siPiezo1 cells did not respond to Yoda1 (Fig.S2A, B), confirming that Yoda1 relies on Piezo1. Therefore, despite triggering similar pERK1/2 levels, Piezo1- and EGF-dependent EGFR internalization lead to different signaling outcomes.

**Figure 4.**
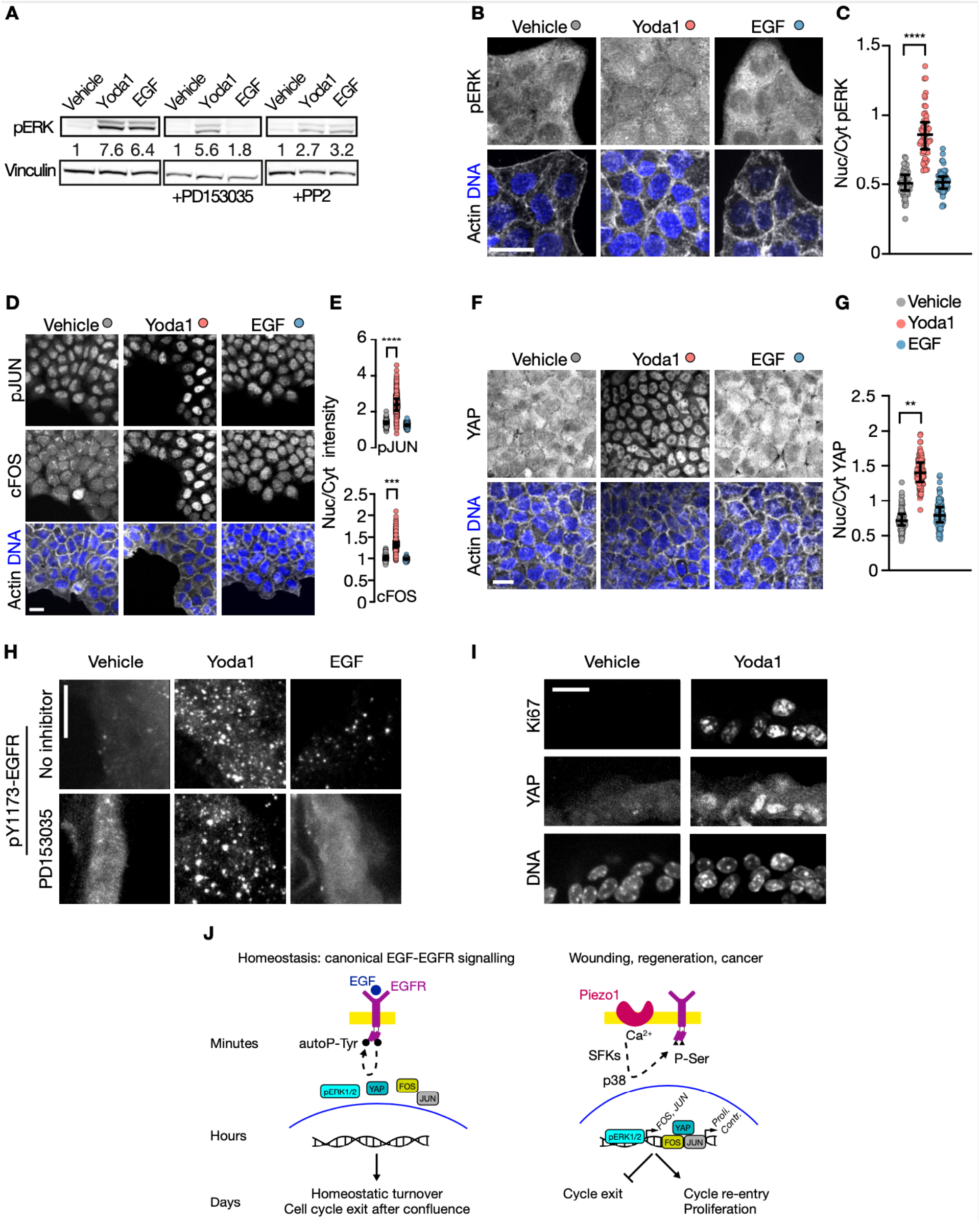
Different downstream signaling in response to Piezo1 activation. **A, B:** Representative western blot (A) and immunostainings (B) of pERK1/2 after 15min of the indicated treatments. Vinculin used as a loading control in A, DNA and Actin shown as spatial references in B. **C**: Quantification of the nuclear vs cytoplasmic enrichment of pERK1/2 signal. **D, F:** Representative immunostainings of pJUN and FOS (D) or YAP (F) after 2h of the indicated treatments. DNA and Actin shown as spatial references. **E, G**: Quantification of the nuclear vs cytoplasmic enrichment of pJUN and FOS (E) or YAP (G) signal. **H, I:** Representative immunostainings of pY1173-EGFR after 15min or of YAP and Ki67 after 48h of the indicated treatments. **J:** Model schematic. Each symbol in C, E, and G represents a cell. Vehicle: 91 (C), 3538 (E), 177 (G); Yoda1: 152 (C), 2543 (E), 301 (G); EGF: 160 (C), 3110 (E), 283 (G). Scale bar= 20μm. Error bars = median±interquartile range. ** p< 0.01, *** p<0.001, **** p< 0.0001 in one-way ANOVA followed by Kruskal-Wallis post-hoc test with Dunn’s correction for multiple comparisons.

Spatiotemporal dysregulation underlies many steps of tumorigenesis (*21, 33*). The duration and strength of nuclear pERK1/2 signaling regulates downstream expression dynamics of the transcription factor FOS, which ultimately drives cell proliferation, differentiation, and transformation (*21, 33, 58*). We found that Yoda1-activation of Piezo1 leads to nuclear accumulation of FOS and phosphorylated JUN (pJUN) within 2h, whereas EGF does not (Fig.4 D, E). Loss of Piezo1 inhibits Yoda1-dependent nuclear translocation of FOS and pJUN (Fig.S2 C&D). FOS and JUN form the dimeric transcription factor AP-1, a key driver of oncogenic growth, migration, and distant organ colonization. In turn, AP-1 cooperates with YAP, which transcriptionally activates dedifferentiation, regeneration, and malignancy, in response to mechanical signals (*5, 7, 59–61*) and Piezo channel activation (*13, 14*). Importantly, Yoda1, but not EGF treatment (Fig.4 F&G), causes nuclear YAP accumulation in a Piezo1-dependent way (Fig.S2 E&F). These results show that Piezo1 activation triggers sustained nuclear accumulation of AP-1 (JUN and FOS) and their mechanosensitive co-activator YAP.

Spatiotemporal dysregulation underlies many steps of tumorigenesis (*21, 33*). The duration and strength of nuclear pERK1/2 signaling regulates downstream expression dynamics of the transcription factor FOS, which ultimately drives cell proliferation, differentiation, and transformation (*21, 33, 58*). We found that Yoda1-activation of Piezo1 leads to nuclear accumulation of FOS and phosphorylated JUN (pJUN) within 2h, whereas EGF does not (Fig.4 D, E). Loss of Piezo1 inhibits Yoda1-dependent nuclear translocation of FOS and pJUN (Fig.S2 C&D). FOS and JUN form the dimeric transcription factor AP-1, a key driver of oncogenic growth, migration, and distant organ colonization. In turn, AP-1 cooperates with YAP, which transcriptionally activates dedifferentiation, regeneration, and malignancy, in response to mechanical signals (*5, 7, 59–61*) and Piezo channel activation (*13, 14*). Importantly, Yoda1, but not EGF treatment (Fig.4 F&G), causes nuclear YAP accumulation in a Piezo1-dependent way (Fig.S2 E&F). These results show that Piezo1 activation triggers sustained nuclear accumulation of AP-1 (JUN and FOS) and their mechanosensitive co-activator YAP.

## Discussion

We show that Piezo1 activates EGFR internalization differently to the canonical EGF-dependent pathway (Fig.4J). During steady state cell turnover, EGF activates EGFR signaling via tyrosine autophosphorylation, receptor internalization, and cytoplasmic ERK activation. Here, we find that acute Piezo1-dependent EGFR activation signals SFK/p38/ nuclear ERK to transcribe early intermediate genes (FOS, JUN) and YAP that promote cell cycle entry. In this way, the same receptor can interpret different inputs to relay separate outcomes, depending on the function of its tyrosine kinase domain. Importantly, we confirmed these findings in and ex vivo mouse lung slice model.

The regulation we found describes a mechanism for one type of paradoxical signaling, where a given signal can trigger opposing outcomes (*62*). Here, EGF ligand binding its receptor contributes to homeostatic cell turnover in epithelia whereas activating the same receptor with stretch drives a proliferative, regenerative outcome, also associated with carcinogenesis. Similar paradoxical systems have been identified to control T cell turnover/death, EGFR recycling/degradation, both depending on soluble ligand concentrations, or autophagy activation or suppression by EGFR depending on its kinase activity and autophosphorylation levels (*37, 50, 63, 64*). Additionally, our previous studies noted a paradoxical signaling response involving Piezo1, depending on the force it experiences: cell extrusion and death upon crowding, or rapid cell division after stretch (*17, 18*). The current study reveals how both EGFR and Piezo1 integrate signaling together to achieve differential outcomes.

SFKs activate p38 in response to radiation and to *Candida* infections to promote survival and infection clearance, respectively, via ligand independent EGFR signaling (Kim et al., 2008; Nikou et al., 2022). Our work identifies mechanical forces via Piezo1 as an endogenous activation mechanism of this axis and reinforce the vision of p38 as a general transactivator of EGFR in response to stressors (antibiotics, anti-cancer drugs, or UV radiation), with a pro-survival and regenerative signaling output different from ligand-induced canonical responses (*47, 48, 54, 55, 65*).

Our findings may be relevant for cancer resistance to anti-EGFR therapy. First, approved EGFR neutralizing antibodies or inhibitors target the receptor’s kinase activity. However, transformation and bad prognosis are associated with increased mechanical signaling and expression of Piezo1, SFKs, p38, and EGFR but not to EGFR phosphorylation status (*13, 16, 27, 66–71*). Second, internalized EGFR has pro-survival functions in cancer cells that are kinase-independent and, thus, unaltered by current tyrosine kinase targeting therapies (*48, 64, 65, 70, 72, 73*). It is tempting to hypothesize that increased mechanical signaling via Piezo1 promotes kinase-independent EGFR functions and drives resistance in cancer cells, as cancer cells experience increased mechanical forces at initiation and migration (*74–76*). Targeting this mechanism could provide an unmet medical need against kinase inhibitor-resistant tumors, e.g. ∼50% of Non-Small Cell Lung Cancers or ∼80% of advanced colorectal cancers or (*70*).

Our findings may also yield practical uses. Given that artificial YAP activation can de-differentiate cells into lineage-restricted stem cells (Panciera et al., 2016) and increase organ size (Camargo et al., 2007), our findings that Yoda1 activates YAP signaling and cell cycle entry *ex vivo* suggest a simple method to obtain tissue-specific stem cells in culture or promote tissue growth.

Our work identifies Piezo1 as a missing link for mechanically activating EGFR and indicates that this activation adopts a different pathway with different outcomes. Mechanical activation of EGFR requires SFK/p38 kinases and serine phosphorylation, rather than EGFR tyrosine phosphorylation, enzymatically separating these two differential outcomes. Given the roles of mechanics in regeneration, oncogenesis, and the increased prevalence of tyrosine kinase inhibitor chemoresistance, our discovery may reveal insight into EGFR signaling during repair and malignancy.

## Materials & Methods

Unless noted, reagents obtained from ThermoFisher.

### Cell culture

Non-verified HeLa, A549, and MDCK-II cells were grown in DMEM (31966021) with 10% FBS (10270106) and 1% Pen/Strep (15070063) in a cell culture incubator at 37ºC with 5% CO_2_. For all experiments, cells were seeded in growth medium. After 24h, growth medium was washed twice with PBS (14190250) and replaced by starvation medium (DMEM+1%Pen/Strep, FBS omitted) for 24h before cell treatment. All cells tested negative for Mycoplasma contamination in periodic PCR tests (Sartorius, 20-700-20).

### siRNA transfection

The day after seeding, 20%-40% confluent cells were transfected with Lipofectamine RNAiMax (13778) and OptiMEM (31985062) following manufacturer’s instructions. siRNA pool (Horizon Discovery, siControl: D-001810-10-50, siPIEZO1: L-020870-03-0050, siCHC: L-004001-01-0020) final concentration was 10nM. siRNA transfection was repeated 24h later. The next day, cells were trypsinized and seeded on glass coverslips in growth medium before starvation overnight and treatment.

### Chemicals and treatments

24h after serum starvation, cells were treated with DMSO (276855), 10µM Yoda1 (Tocris, 5586, reconstituted in DMSO) or 10ng/mL EGF (Peprotech, 400-25, reconstituted in deionized water) for indicated durations. Experiments involving inhibitors included a 30-min pre-treatment with 1µM PD153035 (Sigma, SML0564), 200nM PP2 (Sigma, P0042), or 10 µM SB202190 (Abcam, ab120638), all reconstituted in DMSO. All treatments were prepared recycling starvation medium.

Shear stress was delivered to cells seeded on microfluidic chambers (Ibidi) using a peristaltic pump (Fisher Scientific, 16609762) with a fluid rate of 4mL of starvation medium per minute, shown used to activate Piezo1 (*12*).

### Calcium imaging

Cells grown on glass coverslips were loaded with 4.5 µM of the calcium-sensitive dye Calbryte-520AM (AAT Bioquest, 20651, reconstituted in DMSO) for 30min in the cell culture incubator (37°C, 5% CO2), followed by washing and 30min of additional incubation for AM cleavage and dye equilibration. Coverslips were then mounted on a recording chamber (Warner Instruments, 642420) and intracellular calcium levels were imaged at RT using a 20x air objective, 2×2 binning, and GFP-compatible epifluorescence settings. Fluobrite DMEM (A1896701) supplemented with 20mM HEPES (15630080) was used to prepare all solutions. A peristaltic pump connected to the recording chamber exchanged solutions.

### Western blot

Cells were seeded on 10cm culture plates and starved and treated when 60-70% confluent. After treatment, plates were placed on ice, washed twice with ice-cold PBS, and lysed in 250µL/plate of RIPA buffer (89901) supplemented with EDTA and protease (78430) and phosphatase (Merck, 524625) inhibitor cocktails. After 1min of incubation on ice, plates were scraped and the lysate placed in tubes, followed by 30-min incubation in ice, vortexing every 5 minutes. Lysates were then centrifuged at 13000 rpm for 10 min at 4ºC. The resulting supernatants were placed in new tubes and their protein content quantified with a BCA kit (23225). 50µg grams of protein adjusted to 10 µL in RIPA and dying (B0007) buffers were denatured at 95ºC for 10 min, spun down, supplemented with sample reducing buffer (B0004), and loaded in 4-12% Bis-Tris gels (NW04120BOX). A well with 5 µL of a pre-stained protein standard (LC5925) was used to track protein separation during electrophoresis at 120V for 1.5h in MOPS SDS buffer (B0001). Proteins were then transferred to nitrocellulose membranes (IB23002) with a dry transfer device (IB21001) at 25V for 8min. Membranes were blocked for 1h at RT with 5% BSA (for phosphorylated targets) or with 5% non-fat powder milk. After overnight primary antibody incubation and 3 5min-washings, membranes were incubated at RT for 1h with secondary antibodies followed by 3 additional 5min-washings, 1-min incubation with ECL substrate (32209) and imaging. Band density was later quantified with the Analyze/Gels tool in Fiji (*77*). In some cases, membranes were then stripped and re-blotted following manufacturer’s instructions (Sigma, 2500). 0.1% Tween-20 (Sigma, P137) TBS (TTBS) was used for blocking solutions and washings.

### Antibodies-Western Blot

Primaries: pY1173-EGFR (R&D Systems, AF1095), pY1068-EGFR (CST, 3777S), pS1046/S1047-EGFR (Abcam, ab76300), pERK1/2 (CST, 4370S) all 1/1000 in 5% BSA-TTBS; Vinculin (700062), GAPDH (Abcam, ab8245) 1/2000 in 5% fat free powder milk-TTBS. Incubated overnight at 4ºC inside Falcon tubes with constant rotation.

Secondary: HRP-conjugated secondary anti-Rabbit antibody (65-6120) diluted 1:2500 in 5% non-fat powder milk-TTBS. Incubated 1h at RT.

### Lung slice obtention, treatment, and staining

Mouse lung slices were obtained adapting an established protocol (*78*) approved according to the Animal Scientific Procedures Act. Mice were killed by inhalation of CO_2_ followed by cervical dislocation. After opening the chest cavity, a 20Gx1.25 in canula was inserted in the trachea through a small incision and used to inflate the lungs with 2% low melting agarose (Fisher, BP1360, 2% in HBSS (14025)). After excision and washing in PBS, the lobes were separated and individually embedded in 4% low melting agarose. After casting on ice, a Leica VT1200S vibratome was used to cut 200 µm-thick slices. Slices were washed and incubated overnight in DMEM/F-12 (11320033) with 10% FBS and 1% Pen/Strep in the cell culture incubator. The next morning, slices were individually transferred to a 24-well plate and treated with DMSO or 10 µM Yoda1. After the indicated times, treatments were aspirated, and slices washed thrice with PBS and fixed with 4% PFA overnight at 4ºC. After 2 30-min washings, slices were blocked in 0.1% Triton X-100 and 1% BSA. 1/100 primary and secondary antibodies were consecutively incubated overnight at 4ºC before incubation with 1/1000 DAPI for 20min and mounting in Prolong Gold (P36930). 0.5% Triton X-100 was used for 3 30-min washings between incubations. All solutions prepared in PBS.

### Fixed cell staining

After treatment, cells grown on glass coverslips were fixed in 4% PFA (28908) for 20min at 37°C, permeabilized with 0.5% Triton X-100 (Sigma, X-100) for 5min at RT and stained with primary (overnight, 4°C) and secondary (45min, RT) antibodies. Coverslips were mounted with Fluoromount-G (004958-02). All solutions prepared and washed with PBS.

### Antibodies-Staining

Primaries: pY1173-EGFR (1/250, R&D Systems, AF1095), pS1046/1047-EGFR (1/250, Abcam, ab76300), total EGFR (1/200, antibodies.com, A86603), EEA1 (1/250, BD, 610457), phospho-ERK (1/200, CST, 4370S), YAP (1/200, SCBT, sc-101199), phospho-cJUN (1/200, Abcam, ab32385), cFOS (1/200, SCBT, sc-166940), Ki67 (1/100, Abcam, ab16667).

Secondaries: 1/250 Alexa Fluor (AF)-conjugated 488 Anti-Rabbit (A11008) and 594 Anti-Mouse (A11005), supplemented with 1/500 AF647 Phalloidin (A22287) and 30 μg/ml Hoechst (62249).

For mouse lung slices, 1/100 antibody dilutions were used.

All mixes prepared in 1% BSA (A7906-100G) in PBS. Imaging

Samples were imaged at 1µm-thick Z displacements through 20X and 40X air and 60X oil objectives of a Nikon Eclipse Ti2-E microscope with a Yokogawa CSU-W1 spinning disk system coupled to an Andor DU-888 camera, and a Toptica multi-laser bed. Settings remained unchanged between conditions.

### Image analysis pY1173 internalization

Maximum Intensity Projections were built from raw ND2 files using the *Extract Images* and *Make projections* functions of the *Process images* macro set developed by Christophe Leterrier in Fiji, available at https://github.com/cleterrier/Process_Images. Next, Cell Profiler (*79*) was used to sequentially 1) segment nuclei using the DNA channel, 2) segment cells on the actin channel using nuclei as seeds, 3) generate masks of the cytoplasm after shrinking segmented cells to omit cell boundaries, 4) enhance speckles in the pY1173-EGFR image using a feature size of 10, 5) segment speckles for counting, and 6) export data.

### Nuclear translocation of YAP, cFOS, and pJUN

Background, intra-, and juxta-nuclear ROIs were manually drawn with Fiji using the mid-height image of each Z-stack. The median intensity value of each ROI was measured and exported to a CSV file later used for background subtraction and Nuclear/Cytoplasmic ratio calculation in RStudio or Excel.

### Statistics

All graphs and statistical analyses were done with Prism 9.3.1 (GraphPad Software). Data are presented as median±interquartile range. Given that data distribution was not gaussian, the statistical significance of the differences between treatments was assessed with one-way ANOVA followed by Kruskal-Wallis post-hoc test with Dunn’s correction for multiple comparisons, as suggested by the software. The threshold for statistical significance was p<0.05.

## Acknowledgements

We thank C. Mulas, M. A. Valverde, and F. X. Real for critical reading of the manuscript, F. Fore, and D.C. Bagley for help with mouse work, and E. Ortiz-Zapater, M. Rigau de Llobet, and the members of the Rosenblatt lab for their patience, technical input, and helpful discussions.

C.P.-P. is the recipient of a Long-Term Fellowship (LT000654/2019-L) from the Human Frontier Science Program organization and a Marie Skłodowska-Curie Fellowship (898067) from the European Union’s Horizon 2020 research and innovation program. This research was funded in part by a Wellcome Trust Investigator Award (221908/Z/20/Z), a Cancer Research UK (DRCNPG-May21\100007), an Academy of Medical Sciences Professorship (APR2\1007), and a King’s College London start-up grant to J.R. For open access, the authors have applied a CC BY-NC-ND 4.0 public copyright license to any Author Accepted Manuscript version arising from this submission.

## Resource availability

All data needed to evaluate the conclusions in the paper are present in the paper and/or the Supplementary Materials. Further information and requests for resources and reagents should be directed to and will be fulfilled by the lead contact, Dr. Carlos Pardo-Pastor.

## Author contributions

C.P.-P. designed and performed research, contributed reagents/analytic tools, analyzed data, and wrote the paper; J.R. discussed research and edited the paper.

## Supplementary figures

**Figure S1.**
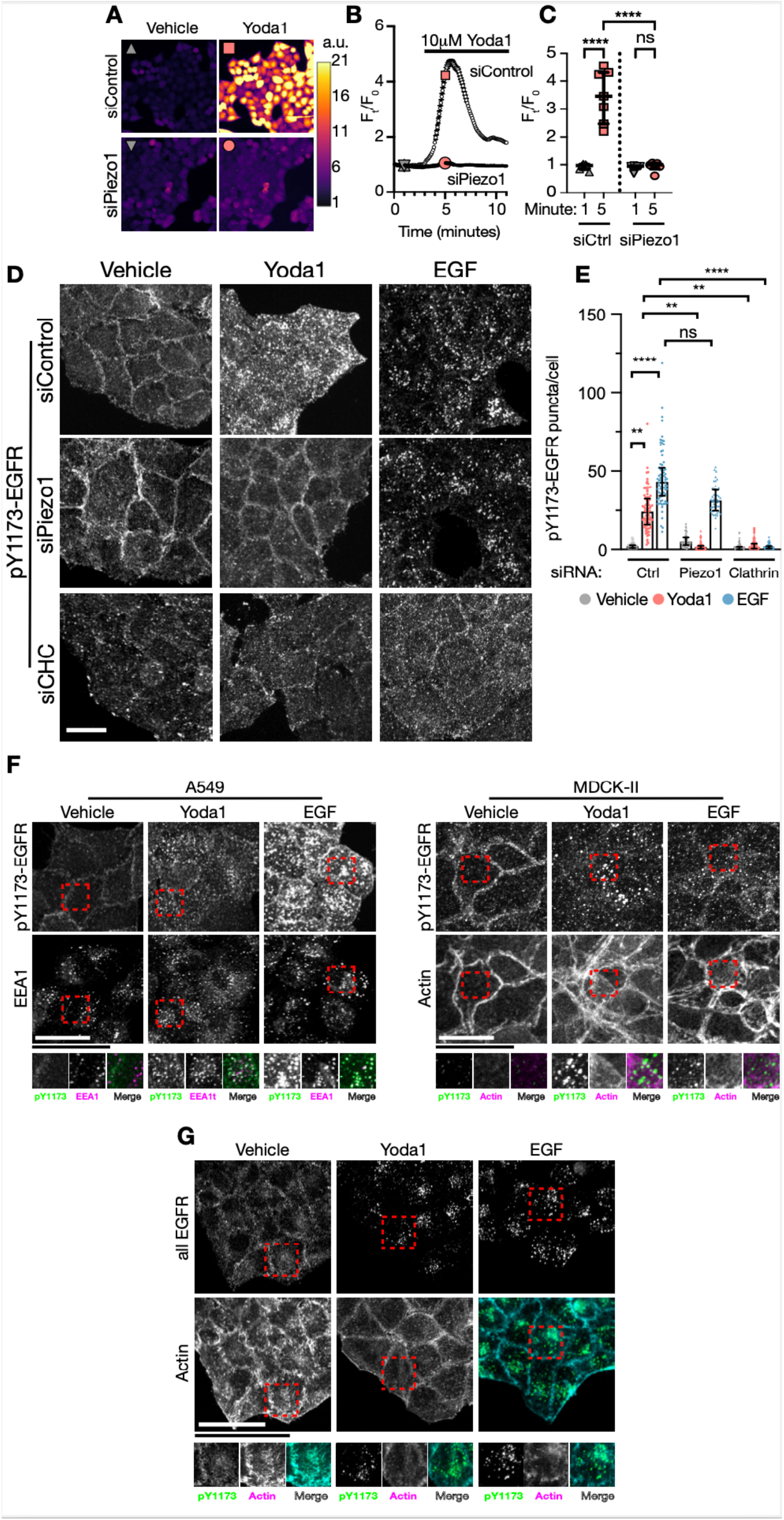
Conserved clathrin mediated EGFR endocytosis in response to Piezo1 activation. **A, B, C:** Representative pseudo-coloured micrographs (A), mean traces (B), and summary statistics (C) of fluorescence intensity in siRNA-transfected HeLa cells loaded with the Ca2+ indicator Calbryte520AM before and after 10μM Yoda1 treatment. siCtrl: 673 cells, 7 independent experiments, 3 transfections; siPiezo1: 1007 cells, 6 independent experiments, 3 transfections. **D:** Representative maximum intensity projections of pY1173-EGFR stainings in control, Piezo1 or CHC KD HeLa cells treated with vehicle, 10μM Yoda1, or 10ng/mL EGF. **E:** Counts of pY1173-EGFR puncta per cell. Each symbol represents a cell. siCtrl (431 Vehicle, 231 Yoda1, 221 EGF), siPiezo1 (611 Vehicle, 702 Yoda1, 298 EGF), siCHC (187 Vehicle, 371 Yoda1, 236 EGF) 3 experiments from 3 siRNA transfections. **F, G:** Representative maximum intensity projections of pY1173-EGFR, EEA1, and Actin stainings in A549 and MDCK-II cells (F) or of total EGFR and Actin in HeLa cells treated for 15min as indicated. Scale bar = 20μm. Error bars = median±interquartile range. ns= non-significant, * p<0.05, **** p< 0.0001, one-way ANOVA followed by Kruskal-Wallis post-hoc test with Dunn’s correction for multiple comparisons.

**Figure S2.**
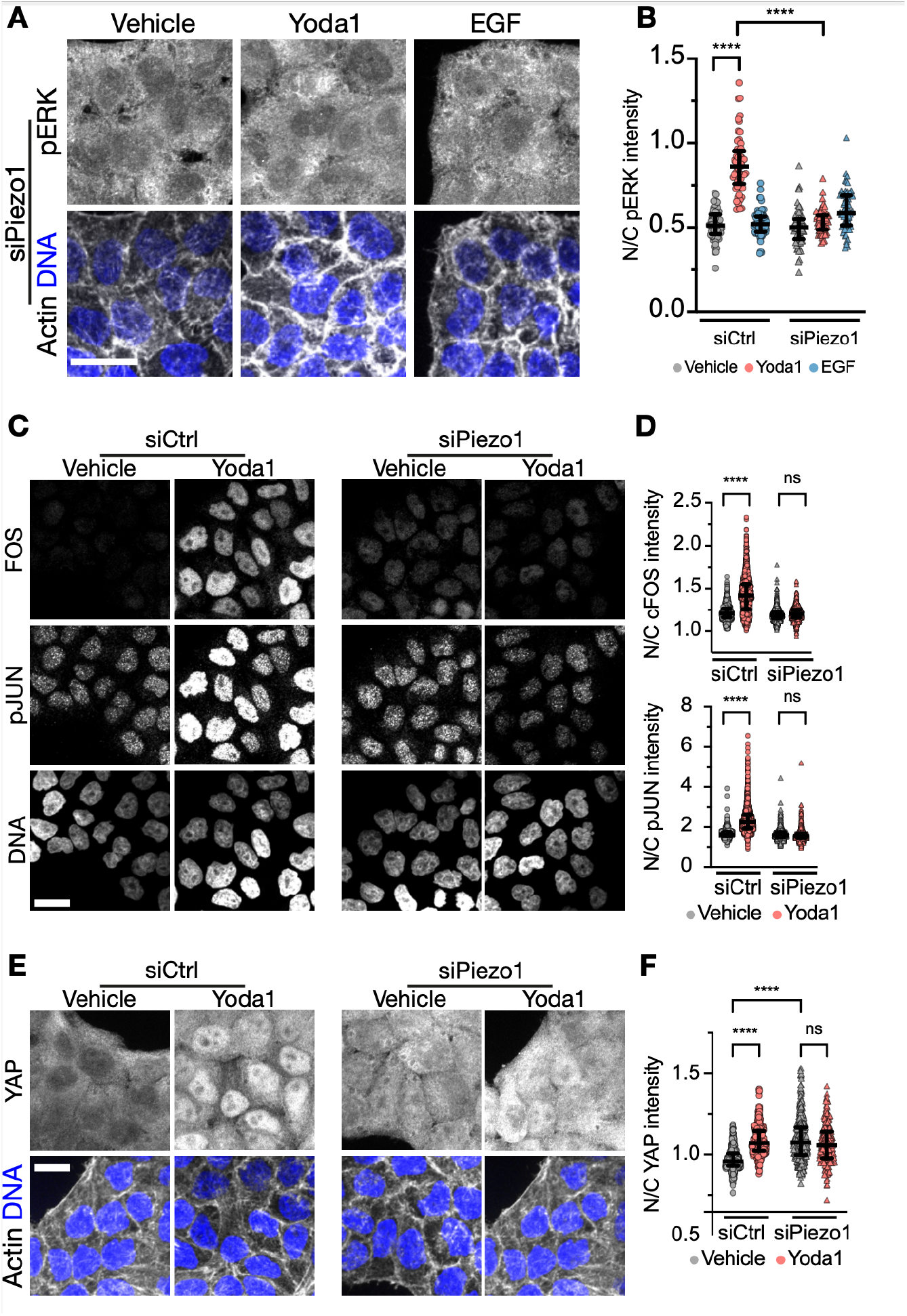
Piezo1 knockdown suppresses nuclear responses to Yoda1. **A, C, E:** Representative immunostainings of pERK1/2 (A) after 15min or FOS, pJUN (C) and YAP (E) after 2h of the indicated treatments. DNA and Actin shown as spatial references. **B, D, F:** Quantification of the nuclear vs cytoplasmic enrichment of pERK1/2 (B), FOS, pJUN (D) and YAP (F) signal. Each symbol in B, D, and F represents a cell. siControl (Vehicle: 91 (C), 11088 (D), 420 (F); Yoda1: 152 (B), 2299 (E), 362 (F); EGF: 160 (C)), siPiezo1 (Vehicle: 146 (B), 1288 (D), 318 (F); Yoda1: 194 (B), 1449 (D), 209 (F). Scale bar= 20μm. Error bars = median±interquartile range. ** p< 0.01, *** p<0.001, **** p< 0.0001 in one-way ANOVA followed by Kruskal-Wallis post-hoc test with Dunn’s correction for multiple comparisons.

## References

1. M. Abercrombie, J. E. Heaysman, Observations on the social behaviour of cells in tissue culture. I. Speed of movement of chick heart fibroblasts in relation to their mutual contacts. Exp. Cell Res. 5, 111–131 (1953).

2. J. Folkman, A. Moscona, Role of cell shape in growth control. Nature. 273, 345–349 (1978).

3. F. M. Watt, P. W. Jordan, C. H. O’Neill, Cell shape controls terminal differentiation of human epidermal keratinocytes. Proc. Natl. Acad. Sci. 85, 5576–5580 (1988).

4. B. Coste, J. Mathur, M. Schmidt, T. J. Earley, S. Ranade, M. J. Petrus, A. E. Dubin, A. Patapoutian, Piezo1 and Piezo2 Are Essential Components of Distinct Mechanically Activated Cation Channels. Science. 330, 55–60 (2010).

5. S. Dupont, L. Morsut, M. Aragona, E. Enzo, S. Giulitti, M. Cordenonsi, F. Zanconato, J. Le Digabel, M. Forcato, S. Bicciato, N. Elvassore, S. Piccolo, Role of YAP/TAZ in mechanotransduction. Nature. 474, 179–183 (2011).

6. R. Latorre, I. Díaz-Franulic, Profile of David Julius and Ardem Patapoutian: 2021 Nobel Laureates in Physiology or Medicine. Proc. Natl. Acad. Sci. 119, e2121015119 (2022).

7. K.-I. Wada, K. Itoga, T. Okano, S. Yonemura, H. Sasaki, Hippo pathway regulation by cell morphology and stress fibers. Development. 138, 3907–3914 (2011).

8. W.-C. Hung, J. R. Yang, C. L. Yankaskas, B. S. Wong, P.-H. Wu, C. Pardo-Pastor, S. A. Serra, M.-J. Chiang, Z. Gu, D. Wirtz, M. A. Valverde, J. T. Yang, J. Zhang, K. Konstantopoulos, Confinement Sensing and Signal Optimization via Piezo1/PKA and Myosin II Pathways. Cell Rep. 15, 1430–1441 (2016).

9. J. Carrillo-Garcia, V. Herrera-Fernández, S. A. Serra, F. Rubio-Moscardo, M. Vogel-Gonzalez, P. Doñate-Macian, C. F. Hevia, C. Pujades, M. A. Valverde, The mechanosensitive Piezo1 channel controls endosome trafficking for an efficient cytokinetic abscission. Sci. Adv. 7, eabi7785 (2021).

10. A. Lai, P. Thurgood, C. D. Cox, C. Chheang, K. Peter, A. Jaworowski, K. Khoshmanesh, S. Baratchi, Piezo1 Response to Shear Stress Is Controlled by the Components of the Extracellular Matrix. ACS Appl. Mater. Interfaces. 14, 40559–40568 (2022).

11. J. Li, B. Hou, S. Tumova, K. Muraki, A. Bruns, M. J. Ludlow, A. Sedo, A. J. Hyman, L. McKeown, R. S. Young, N. Y. Yuldasheva, Y. Majeed, L. A. Wilson, B. Rode, M. A. Bailey, H. R. Kim, Z. Fu, D. A. L. Carter, J. Bilton, H. Imrie, P. Ajuh, T. N. Dear, R. M. Cubbon, M. T. Kearney, K. R. Prasad, P. C. Evans, J. F. X. Ainscough, D. J. Beech, Piezo1 integration of vascular architecture with physiological force. Nature. 515, 279–282 (2014).

12. S. Mylvaganam, J. Plumb, B. Yusuf, R. Li, C.-Y. Lu, L. A. Robinson, S. A. Freeman, S. Grinstein, The spectrin cytoskeleton integrates endothelial mechanoresponses. Nat. Cell Biol. 24, 1226–1238 (2022).

13. C. Pardo-Pastor, F. Rubio-Moscardo, M. Vogel-González, S. A. Serra, A. Afthinos, S. Mrkonjic, O. Destaing, J. F. Abenza, J. M. Fernández-Fernández, X. Trepat, C. Albiges-Rizo, K. Konstantopoulos, M. A. Valverde, Piezo2 channel regulates RhoA and actin cytoskeleton to promote cell mechanobiological responses. Proc. Natl. Acad. Sci. 115, 1925–1930 (2018).

14. M. M. Pathak, J. L. Nourse, T. Tran, J. Hwe, J. Arulmoli, D. T. T. Le, E. Bernardis, L. A. Flanagan, F. Tombola, Stretch-activated ion channel Piezo1 directs lineage choice in human neural stem cells. Proc. Natl. Acad. Sci. 111, 16148–16153 (2014).

15. J. R. Holt, W.-Z. Zeng, E. L. Evans, S.-H. Woo, S. Ma, H. Abuwarda, M. Loud, A. Patapoutian, M. M. Pathak, Spatiotemporal dynamics of PIEZO1 localization controls keratinocyte migration during wound healing. eLife. 10, e65415 (2021).

16. X. Chen, S. Wanggou, A. Bodalia, M. Zhu, W. Dong, J. J. Fan, W. C. Yin, H.-K. Min, M. Hu, D. Draghici, W. Dou, F. Li, F. J. Coutinho, H. Whetstone, M. M. Kushida, P. B. Dirks, Y. Song, C. Hui, Y. Sun, L.-Y. Wang, X. Li, X. Huang, A Feedforward Mechanism Mediated by Mechanosensitive Ion Channel PIEZO1 and Tissue Mechanics Promotes Glioma Aggression. Neuron. 100, 799–815.e7 (2018).

17. G. T. Eisenhoffer, P. D. Loftus, M. Yoshigi, H. Otsuna, C.-B. Chien, P. A. Morcos, J. Rosenblatt, Crowding induces live cell extrusion to maintain homeostatic cell numbers in epithelia. Nature. 484, 546–549 (2012).

18. S. A. Gudipaty, J. Lindblom, P. D. Loftus, M. J. Redd, K. Edes, C. F. Davey, V. Krishnegowda, J. Rosenblatt, Mechanical stretch triggers rapid epithelial cell division through Piezo1. Nature. 543, 118–121 (2017).

19. B. W. Benham-Pyle, B. L. Pruitt, W. J. Nelson, Mechanical strain induces E-cadherin-dependent Yap1 and - catenin activation to drive cell cycle entry. Science. 348, 1024–1027 (2015).

20. Y. Han, C. Liu, D. Zhang, H. Men, L. Huo, Q. Geng, S. Wang, Y. Gao, W. Zhang, Y. Zhang, Z. Jia, Mechanosensitive ion channel Piezo1 promotes prostate cancer development through the activation of the Akt/mTOR pathway and acceleration of cell cycle. Int. J. Oncol. (2019), doi:10.3892/ijo.2019.4839.

21. R. Avraham, Y. Yarden, Feedback regulation of EGFR signalling: decision making by early and delayed loops. Nat. Rev. Mol. Cell Biol. 12, 104–117 (2011).

22. P. E. Farahani, S. B. Lemke, E. Dine, G. Uribe, J. E. Toettcher, C. M. Nelson, Substratum stiffness regulates Erk signaling dynamics through receptor-level control. Cell Rep. 37, 110181 (2021).

23. M. A. Lemmon, J. Schlessinger, Cell Signaling by Receptor Tyrosine Kinases. Cell. 141, 1117–1134 (2010).

24. N. Prenzel, E. Zwick, H. Daub, M. Leserer, R. Abraham, C. Wallasch, A. Ullrich, EGF receptor transactivation by G-protein-coupled receptors requires metalloproteinase cleavage of proHB-EGF. Nature. 402, 884–888 (1999).

25. L. B. Rosen, M. E. Greenberg, Stimulation of growth factor receptor signal transduction by activation of voltage-sensitive calcium channels. Proc. Natl. Acad. Sci. 93, 1113–1118 (1996).

26. S. Sigismund, E. Argenzio, D. Tosoni, E. Cavallaro, S. Polo, P. P. Di Fiore, Clathrin-mediated internalization is essential for sustained EGFR signaling but dispensable for degradation. Dev. Cell. 15, 209–219 (2008).

27. S. Belli, D. Esposito, A. Servetto, A. Pesapane, L. Formisano, R. Bianco, c-Src and EGFR Inhibition in Molecular Cancer Therapy: What Else Can We Improve? Cancers. 12, 1489 (2020).

28. N. Hino, L. Rossetti, A. Marín-Llauradó, K. Aoki, X. Trepat, M. Matsuda, T. Hirashima, ERK-Mediated Mechanochemical Waves Direct Collective Cell Polarization. Dev. Cell. 53, 646–660.e8 (2020).

29. H. Iwasaki, S. Eguchi, H. Ueno, F. Marumo, Y. Hirata, Mechanical stretch stimulates growth of vascular smooth muscle cells via epidermal growth factor receptor. Am. J. Physiol. Heart Circ. Physiol. 278, H521–529 (2000).

30. J.-H. Kim, A. R. Asthagiri, Matrix stiffening sensitizes epithelial cells to EGF and enables the loss of contact inhibition of proliferation. J. Cell Sci. 124, 1280–1287 (2011).

31. S. K. Kuwada, X. Li, Integrin α5/β1 Mediates Fibronectin-dependent Epithelial Cell Proliferation through Epidermal Growth Factor Receptor Activation. Mol. Biol. Cell. 11, 2485–2496 (2000).

32. L. Moro, M. Venturino, C. Bozzo, L. Silengo, F. Altruda, L. Beguinot, G. Tarone, P. Defilippi, Integrins induce activation of EGF receptor: role in MAP kinase induction and adhesion-dependent cell survival. EMBO J. 17, 6622–6632 (1998).

33. L. O. Murphy, J. Blenis, MAPK signal specificity: the right place at the right time. Trends Biochem. Sci. 31, 268–275 (2006).

34. V. Umesh, A. D. Rape, T. A. Ulrich, S. Kumar, Microenvironmental Stiffness Enhances Glioma Cell Proliferation by Stimulating Epidermal Growth Factor Receptor Signaling. PLoS ONE. 9, e101771 (2014).

35. S. Yano, M. Komine, M. Fujimoto, H. Okochi, K. Tamaki, Mechanical Stretching In Vitro Regulates Signal Transduction Pathways and Cellular Proliferation in Human Epidermal Keratinocytes. J. Invest. Dermatol. 122, 783–790 (2004).

36. G. Caldieri, E. Barbieri, G. Nappo, A. Raimondi, M. Bonora, A. Conte, L. G. G. C. Verhoef, S. Confalonieri, M. G. Malabarba, F. Bianchi, A. Cuomo, T. Bonaldi, E. Martini, D. Mazza, P. Pinton, C. Tacchetti, S. Polo, P. P. Di Fiore, S. Sigismund, Reticulon 3–dependent ER-PM contact sites control EGFR nonclathrin endocytosis. Science. 356, 617–624 (2017).

37. S. Sigismund, V. Algisi, G. Nappo, A. Conte, R. Pascolutti, A. Cuomo, T. Bonaldi, E. Argenzio, L. G. G. C. Verhoef, E. Maspero, F. Bianchi, F. Capuani, A. Ciliberto, S. Polo, P. P. Di Fiore, Threshold-controlled ubiquitination of the EGFR directs receptor fate. EMBO J. 32, 2140–2157 (2013).

38. S. Sigismund, T. Woelk, C. Puri, E. Maspero, C. Tacchetti, P. Transidico, P. P. Di Fiore, S. Polo, Clathrin-independent endocytosis of ubiquitinated cargos. Proc. Natl. Acad. Sci. 102, 2760–2765 (2005).

39. D. J. Tschumperlin, G. Dai, I. V. Maly, T. Kikuchi, L. H. Laiho, A. K. McVittie, K. J. Haley, C. M. Lilly, P. T. C. So, D. A. Lauffenburger, R. D. Kamm, J. M. Drazen, Mechanotransduction through growth-factor shedding into the extracellular space. Nature. 429, 83–86 (2004).

40. L. Moro, L. Dolce, S. Cabodi, E. Bergatto, E. Boeri Erba, M. Smeriglio, E. Turco, S. F. Retta, M. G. Giuffrida, M. Venturino, J. Godovac-Zimmermann, A. Conti, E. Schaefer, L. Beguinot, C. Tacchetti, P. Gaggini, L. Silengo, G. Tarone, P. Defilippi, Integrin-induced epidermal growth factor (EGF) receptor activation requires c-Src and p130Cas and leads to phosphorylation of specific EGF receptor tyrosines. J. Biol. Chem. 277, 9405–9414 (2002).

41. M. Saxena, S. Liu, B. Yang, C. Hajal, R. Changede, J. Hu, H. Wolfenson, J. Hone, M. P. Sheetz, EGFR and HER2 activate rigidity sensing only on rigid matrices. Nat. Mater. 16, 775–781 (2017).

42. J. Li, C. Ma, Y. Huang, J. Luo, C. Huang, Differential requirement of EGF receptor and its tyrosine kinase for AP-1 transactivation induced by EGF and TPA. Oncogene. 22, 211–219 (2003).

43. M. P. Oksvold, E. Skarpen, B. Lindeman, N. Roos, H. S. Huitfeldt, Immunocytochemical Localization of Shc and Activated EGF Receptor in Early Endosomes After EGF Stimulation of HeLa Cells. J. Histochem. Cytochem. 48, 21–33 (2000).

44. M. Perez Verdaguer, T. Zhang, J. A. Paulo, S. Gygi, S. C. Watkins, H. Sakurai, A. Sorkin, Mechanism of p38 MAPK–induced EGFR endocytosis and its crosstalk with ligand-induced pathways. J. Cell Biol. 220 (2021), doi:10.1083/jcb.202102005.

45. B. J. McHugh, R. Buttery, Y. Lad, S. Banks, C. Haslett, T. Sethi, Integrin activation by Fam38A uses a novel mechanism of R-Ras targeting to the endoplasmic reticulum. J. Cell Sci. 123, 51–61 (2010).

46. R. Syeda, J. Xu, A. E. Dubin, B. Coste, J. Mathur, T. Huynh, J. Matzen, J. Lao, D. C. Tully, I. H. Engels, H. M. Petrassi, A. M. Schumacher, M. Montal, M. Bandell, A. Patapoutian, Chemical activation of the mechanotransduction channel Piezo1. eLife. 4, e07369 (2015).

47. M. R. Frey, R. S. Dise, K. L. Edelblum, D. B. Polk, p38 kinase regulates epidermal growth factor receptor downregulation and cellular migration. EMBO J. 25, 5683–5692 (2006).

48. T. Tanaka, Y. Zhou, T. Ozawa, R. Okizono, A. Banba, T. Yamamura, E. Oga, A. Muraguchi, H. Sakurai, Ligand-activated epidermal growth factor receptor (EGFR) signaling governs endocytic trafficking of unliganded receptor monomers by non-canonical phosphorylation. J. Biol. Chem. 293, 2288–2301 (2018).

49. Z. Wang, M. F. Moran, Requirement for the Adapter Protein GRB2 in EGF Receptor Endocytosis. Science. 272, 1935–1938 (1996).

50. Y. Wei, Z. Zou, N. Becker, M. Anderson, R. Sumpter, G. Xiao, L. Kinch, P. Koduru, C. S. Christudass, R. W. Veltri, N. V. Grishin, M. Peyton, J. Minna, G. Bhagat, B. Levine, EGFR-Mediated Beclin 1 Phosphorylation in Autophagy Suppression, Tumor Progression, and Tumor Chemoresistance. Cell. 154, 1269–1284 (2013).

51. B. Sullivan, T. Light, V. Vu, A. Kapustka, K. Hristova, D. Leckband, Mechanical disruption of E-cadherin complexes with epidermal growth factor receptor actuates growth factor–dependent signaling. Proc. Natl. Acad. Sci. 119, e2100679119 (2022).

52. N. M. Blythe, K. Muraki, M. J. Ludlow, V. Stylianidis, H. T. J. Gilbert, E. L. Evans, K. Cuthbertson, R. Foster, J. Swift, J. Li, M. J. Drinkhill, F. A. van Nieuwenhoven, K. E. Porter, D. J. Beech, N. A. Turner, Mechanically activated Piezo1 channels of cardiac fibroblasts stimulate p38 mitogen-activated protein kinase activity and interleukin-6 secretion. J. Biol. Chem. 294, 17395–17408 (2019).

53. M. V. Grandal, L. M. Grøvdal, L. Henriksen, M. H. Andersen, M. R. Holst, I. H. Madshus, B. van Deurs, Differential Roles of Grb2 and AP-2 in p38MAPK- and EGF-Induced EGFR Internalization: p38 MAPK and EGFR internalization. Traffic. 13, 576–585 (2012).

54. S. Vergarajauregui, A. S. Miguel, R. Puertollano, Activation of p38 Mitogen-Activated Protein Kinase Promotes Epidermal Growth Factor Receptor Internalization. Traffic. 7, 686–698 (2006).

55. Y. Zwang, Y. Yarden, p38 MAP kinase mediates stress-induced internalization of EGFR: implications for cancer chemotherapy. EMBO J. 25, 4195–4206 (2006).

56. S. Adachi, H. Natsume, J. Yamauchi, R. Matsushima-Nishiwaki, A. K. Joe, H. Moriwaki, O. Kozawa, p38 MAP kinase controls EGF receptor downregulation via phosphorylation at Ser1046/1047. Cancer Lett. 277, 108–113 (2009).

57. P. Wu, P. Wee, J. Jiang, X. Chen, Z. Wang, Differential Regulation of Transcription Factors by Location-Specific EGF Receptor Signaling via a Spatio-Temporal Interplay of ERK Activation. PLOS ONE. 7, e41354 (2012).

58. S. D. M. Santos, P. J. Verveer, P. I. H. Bastiaens, Growth factor-induced MAPK network topology shapes Erk response determining PC-12 cell fate. Nat. Cell Biol. 9, 324–330 (2007).

59. T. Panciera, L. Azzolin, A. Fujimura, D. Di Biagio, C. Frasson, S. Bresolin, S. Soligo, G. Basso, S. Bicciato, A. Rosato, M. Cordenonsi, S. Piccolo, Induction of Expandable Tissue-Specific Stem/Progenitor Cells through Transient Expression of YAP/TAZ. Cell Stem Cell. 19, 725–737 (2016).

60. S. Yui, L. Azzolin, M. Maimets, M. T. Pedersen, R. P. Fordham, S. L. Hansen, H. L. Larsen, J. Guiu, M. R. P. Alves, C. F. Rundsten, J. V. Johansen, Y. Li, C. D. Madsen, T. Nakamura, M. Watanabe, O. H. Nielsen, P. J. Schweiger, S. Piccolo, K. B. Jensen, YAP/TAZ-Dependent Reprogramming of Colonic Epithelium Links ECM Remodeling to Tissue Regeneration. Cell Stem Cell. 22, 35–49.e7 (2018).

61. F. Zanconato, M. Forcato, G. Battilana, L. Azzolin, E. Quaranta, B. Bodega, A. Rosato, S. Bicciato, M. Cordenonsi, S. Piccolo, Genome-wide association between YAP/TAZ/TEAD and AP-1 at enhancers drives oncogenic growth. Nat. Cell Biol. 17, 1218–1227 (2015).

62. Y. Hart, U. Alon, The Utility of Paradoxical Components in Biological Circuits. Mol. Cell. 49, 213–221 (2013).

63. Y. Hart, S. Reich-Zeliger, Y. E. Antebi, I. Zaretsky, A. E. Mayo, U. Alon, N. Friedman, Paradoxical Signaling by a Secreted Molecule Leads to Homeostasis of Cell Levels. Cell. 158, 1022–1032 (2014).

64. X. Tan, N. Thapa, Y. Sun, R. A. Anderson, A Kinase-Independent Role for EGF Receptor in Autophagy Initiation. Cell. 160, 145–160 (2015).

65. A. Tomas, S. O. Vaughan, T. Burgoyne, A. Sorkin, J. A. Hartley, D. Hochhauser, C. E. Futter, WASH and Tsg101/ALIX-dependent diversion of stress-internalized EGFR from the canonical endocytic pathway. Nat. Commun. 6, 7324 (2015).

66. M. E. Fernández-Sánchez, S. Barbier, J. Whitehead, G. Béalle, A. Michel, H. Latorre-Ossa, C. Rey, L. Fouassier, Claperon, L. Brullé, E. Girard, N. Servant, T. Rio-Frio, H. Marie, S. Lesieur, C. Housset, J.-L. Gennisson, M. Tanter, C. Ménager, S. Fre, S. Robine, E. Farge, Mechanical induction of the tumorigenic β-catenin pathway by tumour growth pressure. Nature. 523, 92–95 (2015).

67. R. Mora Vidal, S. Regufe da Mota, A. Hayden, H. Markham, J. Douglas, G. Packham, S. J. Crabb, Urology, in press, doi:10.1016/j.urology.2017.10.041.

68. T. Panciera, A. Citron, D. Di Biagio, G. Battilana, A. Gandin, S. Giulitti, M. Forcato, S. Bicciato, V. Panzetta, S. Fusco, L. Azzolin, A. Totaro, A. P. Dei Tos, M. Fassan, V. Vindigni, F. Bassetto, A. Rosato, G. Brusatin, M. Cordenonsi, S. Piccolo, Reprogramming normal cells into tumour precursors requires ECM stiffness and oncogene-mediated changes of cell mechanical properties. Nat. Mater. (2020), doi:10.1038/s41563-020-0615-x.

69. M. J. Paszek, N. Zahir, K. R. Johnson, J. N. Lakins, G. I. Rozenberg, A. Gefen, C. A. Reinhart-King, S. S. Margulies, M. Dembo, D. Boettiger, D. A. Hammer, V. M. Weaver, Tensional homeostasis and the malignant phenotype. Cancer Cell. 8, 241–254 (2005).

70. R. Thomas, Z. Weihua, Rethink of EGFR in Cancer With Its Kinase Independent Function on Board. Front. Oncol. 9, 800 (2019).

71. W. Zhou, X. Liu, J. W. M. van Wijnbergen, L. Yuan, Y. Liu, C. Zhang, W. Jia, Identification of PIEZO1 as a potential prognostic marker in gliomas. Sci. Rep. 10, 16121 (2020).

72. F. Walker, A. Kato, L. J. Gonez, M. L. Hibbs, N. Pouliot, A. Levitzki, A. W. Burgess, Activation of the Ras/Mitogen-Activated Protein Kinase Pathway by Kinase-Defective Epidermal Growth Factor Receptors Results in Cell Survival but Not Proliferation. Mol. Cell. Biol. 18, 7192–7204 (1998).

73. Z. Weihua, R. Tsan, W.-C. Huang, Q. Wu, C.-H. Chiu, I. J. Fidler, M.-C. Hung, Survival of Cancer Cells Is Maintained by EGFR Independent of Its Kinase Activity. Cancer Cell. 13, 385–393 (2008).

74. F. Broders-Bondon, T. H. Nguyen Ho-Bouldoires, M.-E. Fernandez-Sanchez, E. Farge, Mechanotransduction in tumor progression: The dark side of the force. J. Cell Biol. 217, 1571–1587 (2018).

75. J. M. Northcott, I. S. Dean, J. K. Mouw, V. M. Weaver, Feeling Stress: The Mechanics of Cancer Progression and Aggression. Front. Cell Dev. Biol. 6, 17 (2018).

76. T. Zulueta-Coarasa, J. Fadul, M. Ahmed, J. Rosenblatt, Physical confinement promotes mesenchymal trans-differentiation of invading transformed cells in vivo. iScience. 25, 105330 (2022).

77. J. Schindelin, I. Arganda-Carreras, E. Frise, V. Kaynig, M. Longair, T. Pietzsch, S. Preibisch, C. Rueden, S. Saalfeld, B. Schmid, J.-Y. Tinevez, D. J. White, V. Hartenstein, K. Eliceiri, P. Tomancak, A. Cardona, Fiji: an open-source platform for biological-image analysis. Nat. Methods. 9, 676–682 (2012).

78. K. M. Akram, L. L. Yates, R. Mongey, S. Rothery, D. C. A. Gaboriau, J. Sanderson, M. Hind, M. Griffiths, C. H. Dean, Live imaging of alveologenesis in precision-cut lung slices reveals dynamic epithelial cell behaviour. Nat. Commun. 10, 1178 (2019).

79. C. McQuin, A. Goodman, V. Chernyshev, L. Kamentsky, B. A. Cimini, K. W. Karhohs, M. Doan, L. Ding, S. M. Rafelski, D. Thirstrup, W. Wiegraebe, S. Singh, T. Becker, J. C. Caicedo, A. E. Carpenter, CellProfiler 3.0: Next-generation image processing for biology. PLOS Biol. 16, e2005970 (2018).

